# ACC Sulcal Morphology as a Neurodevelopmental Marker of Proactive and Reactive Cognitive Control among Adolescents

**DOI:** 10.64898/2026.11.16.764094

**Authors:** Marie Lassaigne, Arnaud Cachia, Daniel S. Pine, Nathan A. Fox, Emilio A. Valadez

## Abstract

Behavioral inhibition (BI) is an early temperament associated with higher risk for later anxiety. BI has been linked to variation in cognitive control, particularly proactive control, a crucial component of executive functions which involves monitoring one’s own behavior and adapting that behavior to achieve a particular goal, which may help explain BI’s association with elevated anxiety risk. Yet little is known about the neurobiology of cognitive control development among children with BI. Variation in anterior cingulate cortex (ACC) sulcal pattern asymmetry, established prenatally and stable across development, is associated with individual differences in cognitive control. This preliminary study examined whether ACC sulcal symmetry is associated with BI, proactive and reactive control (measured via the AX-CPT) in adolescents. Structural T1-weighted MRI was acquired from 62 adolescents aged 12 to 15 years. ACC patterns in each hemisphere were classified as ‘single type’ or ‘double parallel type’ based on the presence of a paracingulate sulcus. Adolescents with symmetric ACC patterns showed lower proactive control (*d’* context; *η²p* = .091) and greater reactive control (BX probe interference effect; *η²p* = .119). These associations held even when tested within the same statistical model and were comparable in magnitude. ACC symmetry was not associated with BI, nor did it interact with BI to predict cognitive control scores. Overall, the findings provide preliminary evidence that ACC sulcal morphology may be a structural marker of variability in cognitive control, with relevance to BI-related neurocognitive risk.

## 1. Introduction

Behavioral inhibition (BI) is an early-emerging temperament characterized by fear or avoidance of novel experiences, people and places (Kagan et al., 1984; Fox et al., 2023). Across longitudinal studies, approximately 40-60% of children with the phenotype go on to develop anxiety disorders, specifically, social anxiety (Clauss & Blackford, 2012; Fox et al., 2023). Thus, about half of children with BI as toddlers do not develop anxious disorders as children or adolescents. Understanding what differentiates those who do from those who do not can inform early identification efforts and novel intervention development.

Multiple studies have identified cognitive control as a key moderator explaining variability in anxiety outcomes among children with BI. Cognitive control allows individuals to develop flexibility in their ability to respond adaptively to a wide variety of experiences (Cachia et al., 2014). The Detection and Dual Control (DDC) framework (Fox et al., 2021, 2023; Valadez et al., 2022), an extension of the dual mechanisms of control theory (Braver, 2012), distinguishes risk in BI related to two forms of cognitive control: automatic and planful control. Whereas automatic control (e.g., inhibitory control, reactive control) relates to high anxiety risk among youth with a history of BI (Drexler et al., 2024; Troller-Renfree et al., 2019b; Valadez et al., 2021, 2022), planful control (e.g., proactive control, task switching) relates to lower anxiety risk for these children (Troller-Renfree et al., 2019a; Valadez et al., 2021, 2022).

Despite evidence for cognitive control as a moderator of the BI-anxiety association, little is known about the neurobiological underpinnings of cognitive control development in the context of BI. Most work linking cognitive control to anxiety risk in behaviorally inhibited youth has relied on task performance or functional measures, which are themselves shaped by experience and maturation and reflect the consequences of a developmental trajectory as much as its origins (Fox et al., 2023; Valadez et al., 2022, 2025). Structural measures depend less on the task or on momentary state, but they also change across development. BI in early childhood predicts thinner dorsal ACC in adulthood (Sylvester et al., 2016), but whether thinner cortex precedes BI or is a result of it is unknown. Cortical sulcal patterns, qualitative features of cortical folding topology and spatial organization, such as the presence or absence of a sulcus, are largely established during fetal development and remain relatively stable across the lifespan (Cachia et al., 2016), unlike cortical thickness. They may therefore index early neurodevelopmental processes that precede, and potentially constrain, subsequent experience-dependent plasticity (Cachia et al., 2021). The ACC is a strong candidate because folding emerges between 10- and 15-weeks’ gestation, shows substantial interindividual variability and is associated with individual differences in executive functioning (Amiez et al., 2018; Cachia et al., 2021).

The ACC is a core region of the cognitive control network. According to the conflict-monitoring hypothesis, the dorsal ACC signals conflict in information processing to the cognitive control system supported through dorsolateral prefrontal cortices (Botvinick, 2007; Botvinick et al., 2001). ACC sulcal morphology falls into two patterns: a ‘single type’, in which only a cingulate sulcus is present, and a ‘double parallel type’, in which a paracingulate sulcus (PCS) runs parallel to the cingulate sulcus (Ono et al., 1990). Asymmetry between hemispheres has been linked to cognitive control across age groups, including ERP markers of conflict monitoring (Huster et al., 2014) and cognitive control efficiency across the lifespan (Del Maschio et al., 2019; Fornito et al., 2004). ACC asymmetry explained about 21% of the variance in Stroop interference in preschoolers (Cachia et al., 2014), and 13.7% in children and adults (Tissier et al., 2018). Brain-behavior associations are generally modest, however, in over 11,000 children, MRI predicted about 20% of the variance in cognition (Pat et al., 2022). Cognition remains the domain best suited for brain-based prediction (Wang et al., 2025), which is why the present study focuses on cognitive control rather than temperament or anxiety directly.

Despite these connections, no study has examined ACC sulcal morphology in relation to proactive and reactive control specifically, nor in a sample characterized by BI and early temperament variation. Prior work has relied on Stroop and Flanker paradigms, which can index cognitive control efficiency but do not dissociate the control modes described by the DDC framework. The AX-CPT allows this dissociation within a single task: d’ context provides a theoretically precise index of proactive control, while the BX probe interference effect indexes reactive control. D’ context reflects the extent to which participants adjust their responses to the cue context, maintaining the A cue to prepare the correct response to the next probe, so that higher scores indicate more effective use of contextual information to guide behavior in advance.

The BX probe interference effect reflects the extent to which participants respond to the probe even when it is not informative, indexing greater difficulty overriding the target response on BX trials and greater reliance on reactive control. Examining both is essential, because the DDC framework predicts that proactive and reactive control relate to anxiety risk in opposite directions, and it is therefore unclear whether an early structural marker of cognitive control would relate to both modes or preferentially to one. Additionally, whether BI or infant reactivity, a precursor to BI, moderates the morphology-cognition relationship remains unknown. The present study addresses these gaps in a sample of 62 adolescents drawn from a longitudinal study of temperament.

### Hypotheses

In this context, we predicted that ACC sulcal symmetry would be associated with lower d’ context scores, replicating the broader cognitive control literature. Given that the AX-CPT also yields an index of reactive control, we examined whether ACC sulcal symmetry was associated with the BX probe interference effect and whether the magnitude of the association differed across the two control modes. Since our sample was recruited for their behaviorally inhibited temperament, we also tested whether ACC symmetry was associated with BI and whether BI moderated the morphology-cognition relationship. Whether infant reactivity moderated these associations was reported in Section S3.

## 2. Materials and methods

### 2.1. Participants

A total of 71 participants aged 12-15 years (*M* = 13.16, *SD* = 1.47) were drawn from the larger longitudinal Temperament over Time Study (TOTS; *N* = 291). Participants underwent structural MRI, cognitive assessments, and observation-based assessments in the laboratory. Of the 71 participants scanned, 5 were excluded due to unusable structural MRI data that prevented sulcal classification (see Section 2.3), resulting in 66 participants with usable MRI data. The study protocols were approved by the Institutional Review Boards of the National Institute of Mental Health Intramural Research Program (MRI laboratory visits) and the University of Maryland, College Park (all other data collection).

### 2.2 MRI Acquisition and Preprocessing

T1-weighted whole-brain structural MRI images were acquired in a 3-Tesla MR750 GE scanner with a 32-channel head coil. The T1-weighted scan was a magnetization-prepared rapid acquisition gradient-echo sequence (MPRAGE; sagittal acquisition; TI/TE = 425/min; flip angle = 7°; FOV = 25.6; Matrix 256 × 256; Slice thickness = 1 mm; bandwidth = 25 Hz).

Raw T1-weighted MRI scans were processed using BrainVISA (version 6.12; http://brainvisa.info/) with standard parameters, which was used for cortical surface reconstruction and sulcal segmentation via the Morphologist toolbox. This approach has been widely used for ACC sulcal classification in previous studies (e.g. Cachia et al., 2016; Gay et al., 2017; Tissier et al., 2018; Wu et al., 2025), as it provides a robust three-dimensional representation of cingulate sulcus and PCS. During automated preprocessing, T1-weighted MRI scans were skull-stripped, and brain tissues were segmented. Spatial normalization was intentionally omitted to avoid biases that could result from sulcal shape distortions introduced by warping. Cortical folds were automatically delineated from the skeleton of the gray matter– cerebrospinal fluid mask; folds were defined as the crevasse bottoms of the cortical “landscape”, where height corresponds to MRI intensity. This method provides a stable and robust representation of sulcal surfaces that is not affected by variations in cortical thickness or gray/white matter contrast (Mangin et al., 2004).

### 2.3 ACC Sulcal Pattern Classification

The sulcal pattern of the dorsal ACC was then classified through visual inspection of the resulting 3D cortical surface reconstructions, following Ono’s classification (Ono et al. 1990).

Each hemisphere was assigned one of two anatomical variants: “single type,” defined by the presence of the cingulate sulcus alone, or “double parallel type,” defined by the additional presence of the PCS. The PCS, when present, was located dorsally to the cingulate sulcus and ran parallel to it (Paus et al. 1996). To disambiguate the confluence of the PCS with the superior rostral sulcus, we defined the anterior border of the PCS as the point where the sulcus extended posteriorly beyond an imaginary vertical line perpendicular to the axis connecting the anterior and posterior commissures (AC-PC) (Yucel et al. 2001). The PCS was considered present only if it extended at least 20 mm horizontally beyond this imaginary vertical line. Patterns were classified as “symmetric” when identical across hemispheres and “asymmetric” when they differed. All labeling was performed by a single rater (M.L.) blind to all participant demographics and scores. Classifications were subsequently reviewed by an expert (A.C.), who corrected any cases deemed mislabeled and who was similarly blinded.

**Figure 1.**
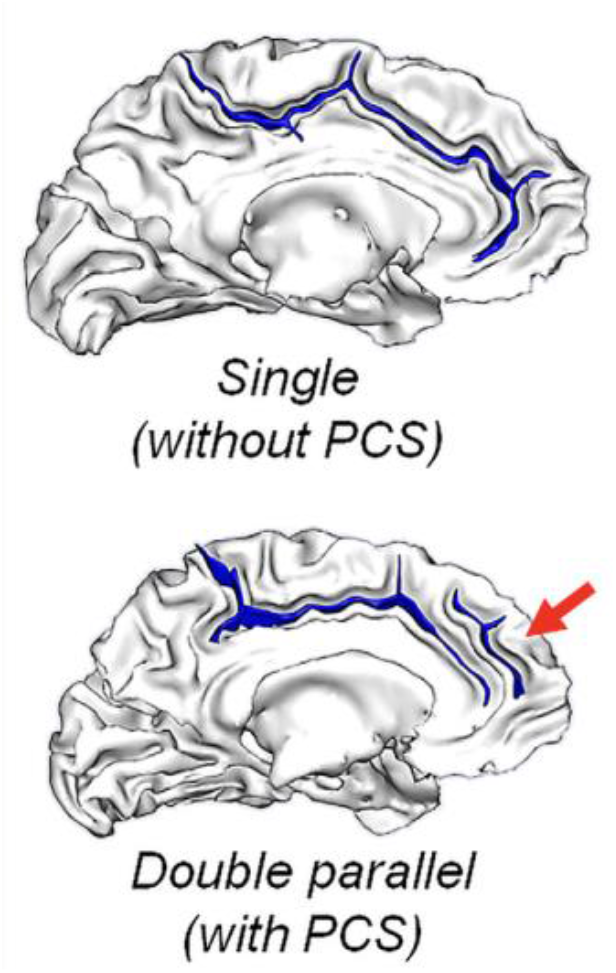
ACC sulcal patterns. Single type = cingulate sulcus only; double parallel type = added paracingulate sulcus (PCS, red arrow). Hemispheres classified as single/double, participants as symmetric or asymmetric. Adapted from Cachia et al. (2021).

### 2.4 Cognitive Control Assessment

Proactive and reactive control were assessed at the 15-year assessment using the AX-Continuous Performance Test (AX-CPT; Cohen et al., 1999). The task presents a continuous series of letter pairs: a cue followed by a probe. Participants were instructed to press a target button only when an “A” cue was followed by an “X” probe, and a nontarget button for all other combinations. The task is dissociable into four trial types: AX (target cue, target probe), AY (target cue, nontarget probe), BX (nontarget cue, target probe), and BY (nontarget cue, nontarget probe). Accurate performance depends on maintaining cue identity in working memory across the cue-probe delay (proactive control) while also responding correctly to differences in the probe letter (reactive control).

In addition to measuring proactive control via d’ context, reactive control was indexed from the same task using the BX probe interference effect, which reflects the extent to which responding is driven by the probe rather than the maintained cue. Two versions were computed by contrasting the BX and BY trials: an error-rate measure and a reaction-time measure, with higher values on each reflecting greater interference by the probe. The error-rate index served as the primary reactive measure and the reaction-time index as a secondary measure.

Trial type proportions favored AX trials (55%), establishing the expectancy on which proactive control depends. Full task parameters and stimulus details are reported in the Supplementary Materials (Section S1). Proactive control was measured via d’ context, a signal detection theory index computed by comparing hit rates on AX trials to false alarm rates on BX trials, with higher scores indicating greater proactive control use (Cohen et al., 1999; Servan-Schreiber et al., 1996).

Participants were required to achieve at least 60% accuracy on BY trials for inclusion in analyses to ensure that task instructions were understood. Of the 66 participants with valid ACC sulcal reconstruction, 3 did not complete the 15-year AX-CPT and 1 completed fewer than one task block yielding a final analytic sample of 62. Trial-level exclusion criteria are described in Section S1.

### 2.5 Behavioral Inhibition and Infant Reactivity

Behavioral inhibition was measured across 2 and 3 years of age. Participants were presented with unfamiliar adults and objects. Behavioral coding was based on toddlers’ approach or avoidance to the novel stimuli and was averaged across the 2- and 3-year assessments, as described in previous reports (Fox et al., 2001; Calkins et al., 1996). In this sample, behavioral inhibition scores were approximately evenly distributed around the mean (*M* = 0.03, *SD* = 0.71, range= −1.64 to 1.50; *n* = 65).

Infant reactivity, a precursor to BI, was measured at age 4 months. Participants were observed while reacting to new objects and new sounds using the standardized Kagan assessment (Kagan & Snidman, 1991). Infant reactivity was categorized as control if showing little emotion or movement, negative reactivity if they became fussy or positive reactivity if they seemed excited. In our neuroimaging subsample, 29 participants showed negative reactivity, 17 positive reactivity, 13 were classified as control and 3 had missing variables.

### 2.6 Statistical Analyses

We first tested whether ACC sulcal symmetry predicted proactive control (d’ context) on the AX-CPT. Then, given the established links between BI and cognitive control, we tested whether ACC sulcal symmetry was associated with BI, and whether BI moderated the association between symmetry and cognitive control. To investigate evidence of the association beyond conventional null hypothesis testing, Bayes factors were computed for the behavioral inhibition analyses (Section S5). All analyses were based on linear models, fitted using R (version 4.3.1) using the “lm” and “Anova” functions within the “car” package. Linear models adjusted for the same covariates: gender, race/ethnicity, and age at scan. Because the sample was predominantly White, with very small cell sizes for other racial/ethnic groups, race/ethnicity was coded as binary with White versus non-White (based on maternal report). To examine whether early temperament moderated the association between ACC morphology and cognitive control, analyses included an ACC symmetry by 4-months reactivity group interaction are reported in Section S3.

We additionally tested whether ACC symmetry predicted reactive control (BX probe interference) and modeled proactive and reactive control jointly to compare the magnitude of the symmetry effect across control modes. A joint model (Section S4) was fit in which d’ context and BX interference were regressed simultaneously on ACC symmetry and covariates (gender, age at scan, behavioral inhibition, race/ethnicity, and reactivity group) using structural equation modeling (SEM, “lavaan” package) with full-information maximum likelihood (FIML). Because the two outcomes are scored in opposite directions (higher d’ context indicates stronger proactive control and higher BX probe interference indicates greater use of reactive control), the difference in the ACC symmetry effect across the two outcomes was tested as the sum of the two standardized coefficients.

### 3.1 Sample Characteristics and ACC Distribution

The final analytic sample comprised 62 adolescents (29 males, 30 females, 3 missing; *M* age = 13.16 years, *SD* = 1.47, range = 12-15) drawn from the TOTS sample.

Of the 62 participants with complete MRI and d’ context data, 38 (61.3%) showed a symmetric ACC sulcal pattern and 24 (38.7%) showed an asymmetric pattern. In the left hemisphere, 44 participants (71.0%) had a single-type pattern and 18 (29.0%) had a double parallel pattern. In the right hemisphere, 44 (71.0%) had a single-type pattern and 18 (29.0%) had a double parallel pattern.

In comparisons to prior studies in healthy samples, the proportion of asymmetric patterns in our sample (38.7%) is broadly consistent. Yucel et al. (2001) reported 44% asymmetric in 171 adults, Cachia et al. (2014) reported 42% in preschoolers, Borst et al. (2014) reported 39% in children and Santacroce et al. (2026) reported 48.5% in 390 young adults from the Human Connectome Project. Tissier et al. (2018) reported a significantly higher proportion (76%), though this is acknowledged in the paper as likely being due to small sample size. The leftward bias in double parallel pattern prevalence observed in the literature (Paus et al., 1996) was not strongly evident in our sample, likely reflecting methodological and sample differences across studies rather than developmental change since the pattern is stable across time. Sample characteristics are presented in Table 1.

**Table 1.**
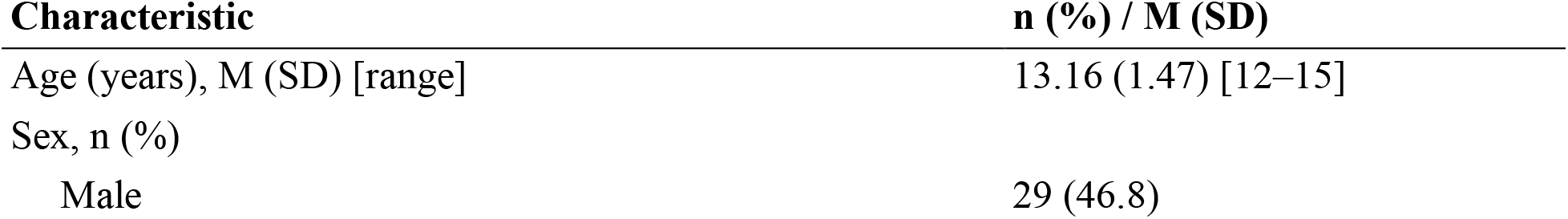
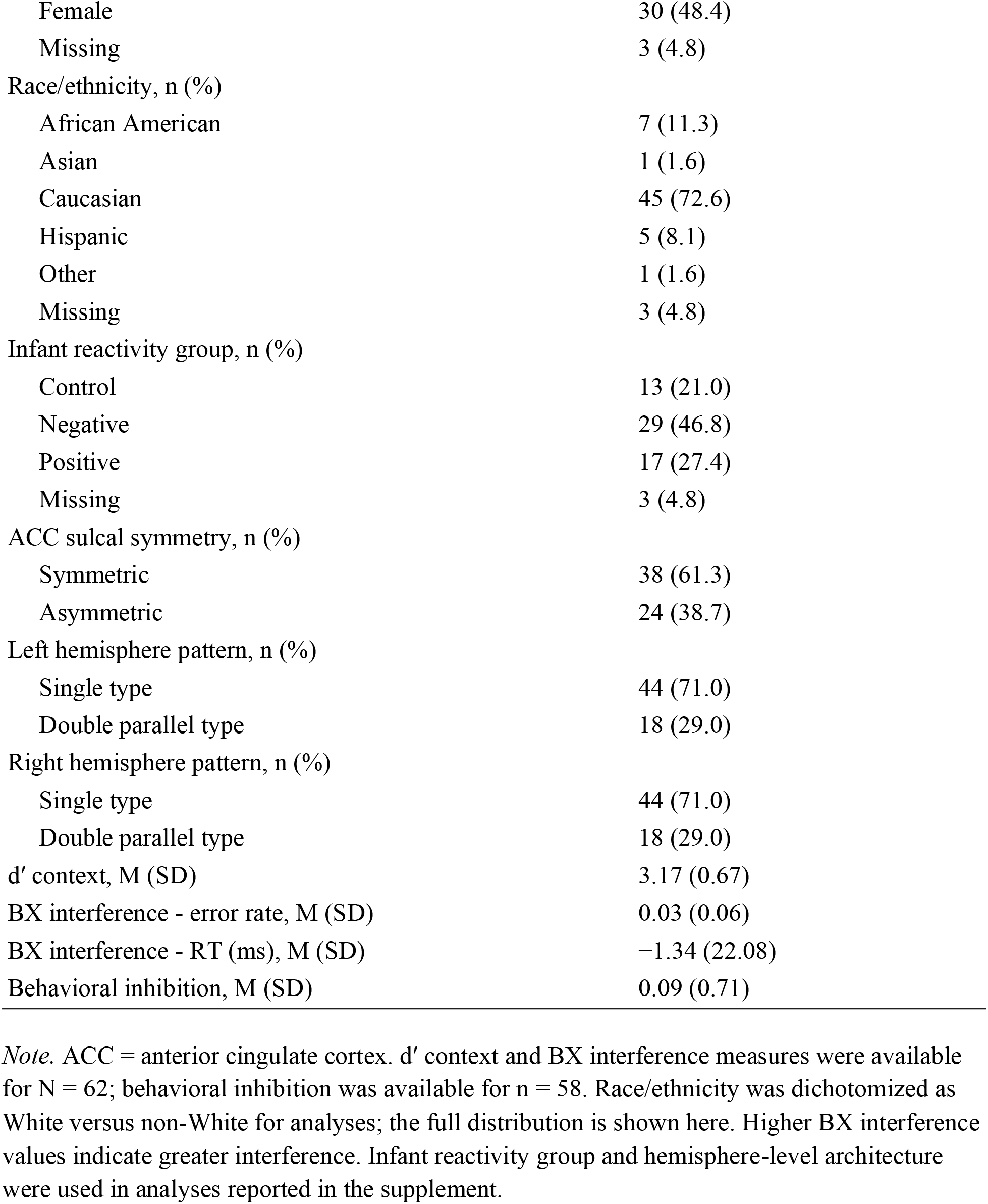
Demographic and Descriptive Characteristics of the Analytic Sample (N = 62)

| **Characteristic** | **n (%) / M (SD)** |
|---|---|
| Age (years), M (SD) [range] | 13.16 (1.47) [12–15] |
| Sex, n (%) |  |
| Male | 29 (46.8) |
| Female | 30 (48.4) |
| Missing | 3 (4.8) |
| Race/ethnicity, n (%) |  |
| African American | 7 (11.3) |
| Asian | 1 (1.6) |
| Caucasian | 45 (72.6) |
| Hispanic | 5 (8.1) |
| Other | 1 (1.6) |
| Missing | 3 (4.8) |
| Infant reactivity group, n (%) |  |
| Control | 13 (21.0) |
| Negative | 29 (46.8) |
| Positive | 17 (27.4) |
| Missing | 3 (4.8) |
| ACC sulcal symmetry, n (%) |  |
| Symmetric | 38 (61.3) |
| Asymmetric | 24 (38.7) |
| Left hemisphere pattern, n (%) |  |
| Single type | 44 (71.0) |
| Double parallel type | 18 (29.0) |
| Right hemisphere pattern, n (%) |  |
| Single type | 44 (71.0) |
| Double parallel type | 18 (29.0) |
| d′ context, M (SD) | 3.17 (0.67) |
| BX interference - error rate, M (SD) | 0.03 (0.06) |
| BX interference - RT (ms), M (SD) | −1.34 (22.08) |
| Behavioral inhibition, M (SD) | 0.09 (0.71) |

*Note.* ACC = anterior cingulate cortex. d′ context and BX interference measures were available for N = 62; behavioral inhibition was available for n = 58. Race/ethnicity was dichotomized as White versus non-White for analyses; the full distribution is shown here. Higher BX interference values indicate greater interference. Infant reactivity group and hemisphere-level architecture were used in analyses reported in the supplement.

### 3.2 ACC Symmetry and Cognitive Control

Descriptive statistics and correlations among the main study variables are presented in Table 2. As expected, ACC sulcal symmetry was significantly associated with 15-year AX-CPT proactive control performance (*F*(1, 54) = 5.42, *p* = .024, *η²p* = .091, *N* = 59), such that adolescents with symmetric ACC sulcal patterns showed lower proactive control (d’ context scores) than those with asymmetric patterns.

**Table 2.** Means, Standard Deviations, and Intercorrelations Among Main Study Variables.

| Variable | M | SD | 1 | 2 | 3 | 4 | 5 | 6 | 7 |
|---|---|---|---|---|---|---|---|---|---|
| 1. ACC symmetry | 0.61 | 0.49 | — |  |  |  |  |  |  |
| 2. d′ context | 3.17 | 0.67 | −.25* | — |  |  |  |  |  |
| 3. BX interference (error) | 0.03 | 0.06 | .30* | −.62** | — |  |  |  |  |
| 4. BX interference (RT) | −1.34 | 22.08 | .21 | −.15 | .35** | — |  |  |  |
| 5. Behavioral inhibition | 0.09 | 0.71 | −.10 | .02 | −.25 | −.07 | — |  |  |
| 6. Age | 13.16 | 1.47 | .09 | .06 | −.11 | −.06 | .03 | — |  |
| 7. Sex | 0.51 | 0.50 | −.20 | −.04 | .03 | −.03 | .30* | .12 | — |

*Note.* N = 62; correlations involving behavioral inhibition are based on n = 58 (pairwise). ACC symmetry coded 0 = asymmetric, 1 = symmetric; Sex coded 0 = male, 1 = female. Higher BX interference values indicate greater interference.
\* p < .05. \*\* p < .01.

**Figure 2.**
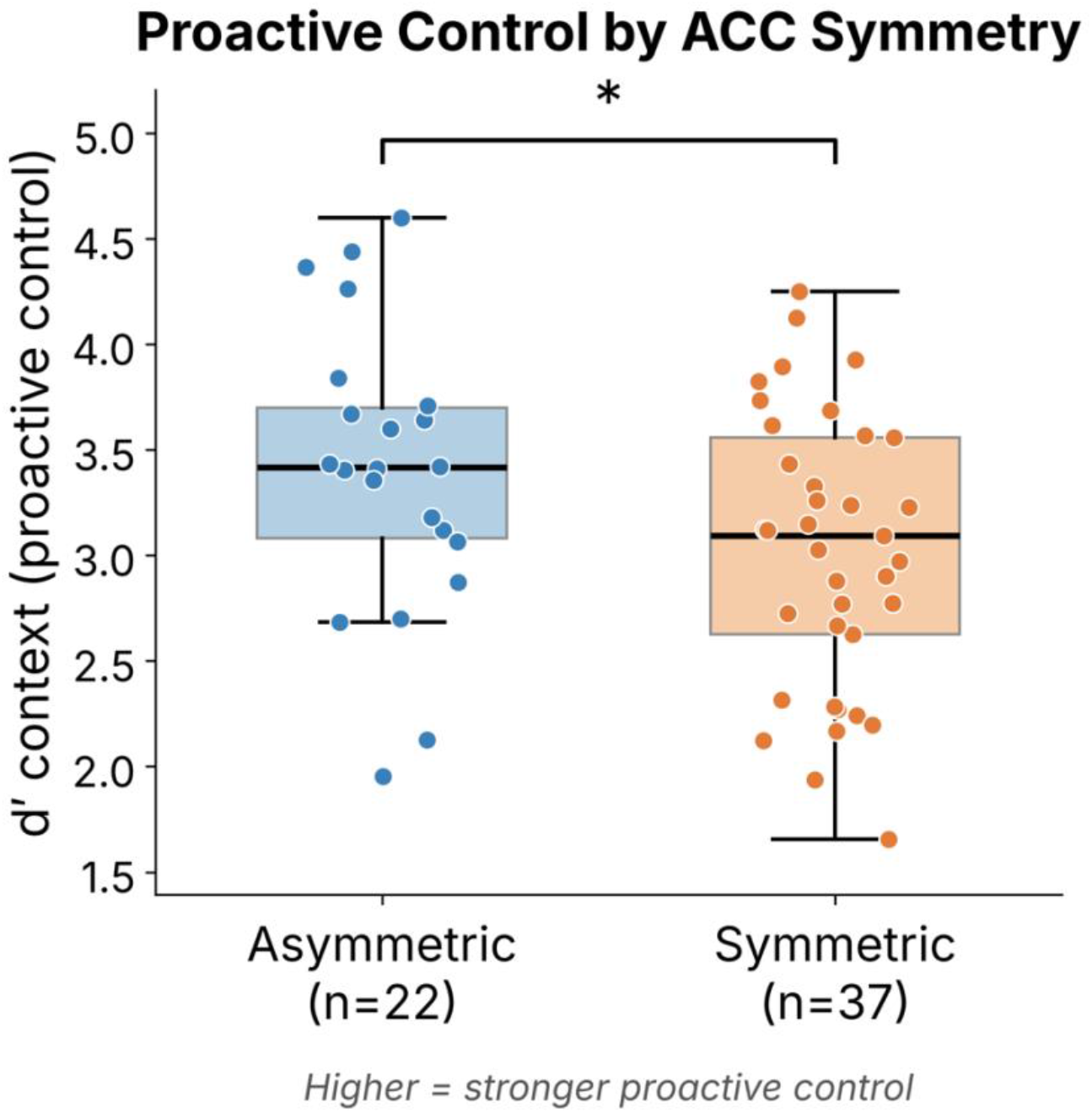
Proactive control (d’ context) by ACC sulcal pattern. Adolescents with asymmetric patterns showed higher d’ context. * p < .05. ** p < .01.

ACC sulcal symmetry was also associated with the BX probe interference error rate (*F*(1, 54) = 7.31, *p* = .009, *η²p* = .119, *N* = 59), such that adolescents with symmetric patterns showed greater reactive control (i.e., greater probe interference) than those with asymmetric patterns. The pattern was in the same direction for the reaction-time measure at a trend level (*F*(1, 54) = 3.96, *p* = .052, *η²p* = .068, *N* = 59).

**Figure 3.**
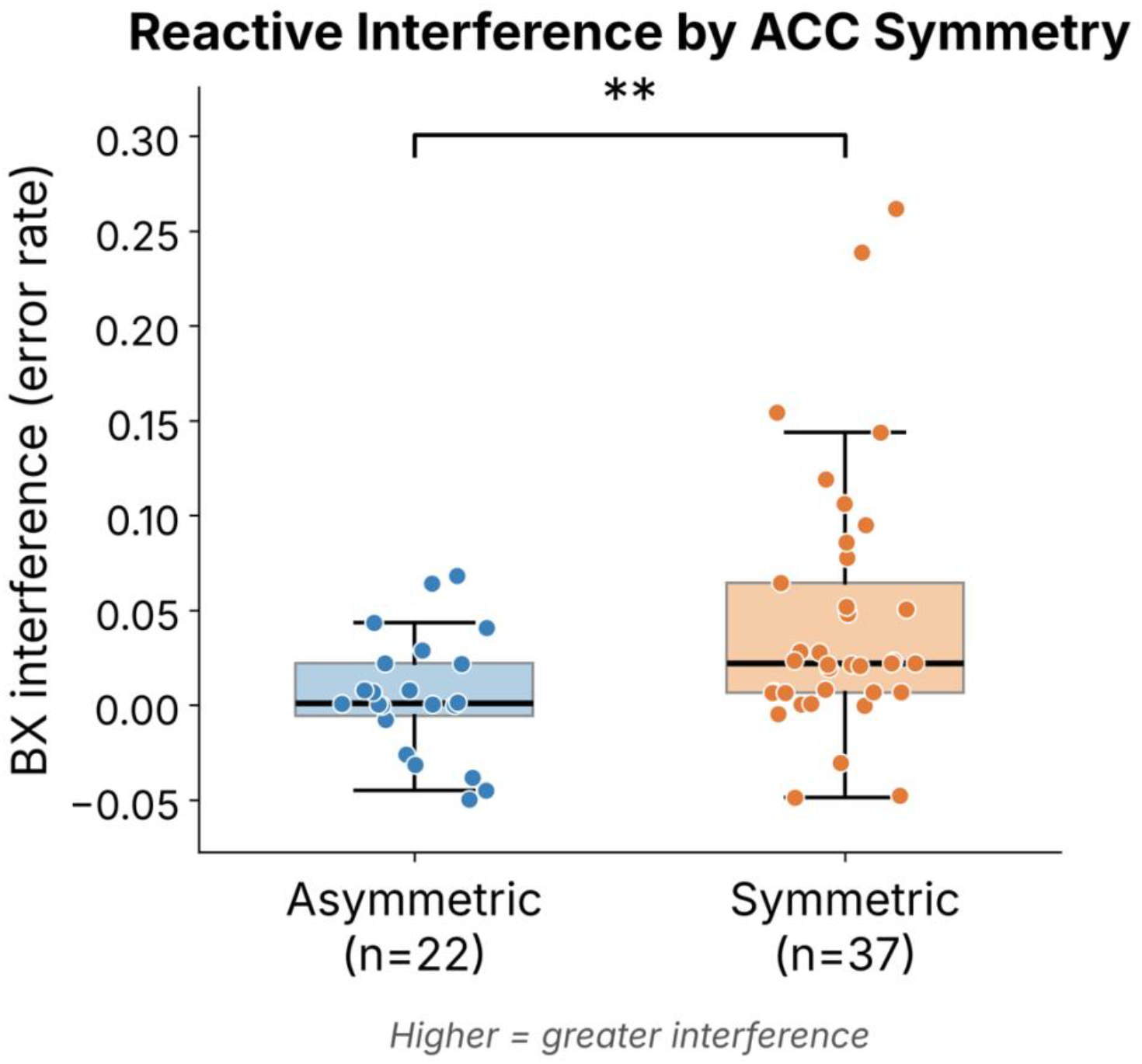
Reactive control (BX interference) by ACC sulcal pattern. Adolescents with asymmetric patterns showed lower interference. * p < .05. ** p < .01.

### 3.3 ACC Symmetry and Behavioral Inhibition

Given the established links between BI and cognitive control, we tested whether ACC sulcal symmetry was associated with BI. ACC sulcal symmetry did not predict BI (*F*(1, 57) = 0.03, *p* = .861, *N* = 62). We further tested whether BI moderated the association between ACC symmetry and cognitive control. The symmetry-by-BI interaction was not significant for proactive control (*F*(1, 51) = 0.44, *p* = .511) or for reactive control (BX interference error rate: *F*(1, 51) = 0.50, *p* = .483). Bayes factors indicated moderate evidence (BF01 = 3.79), whereas the symmetry-by-BI interactions only provided weak evidence supporting no association between ACC sulcal symmetry and BI (proactive control: BF01 = 2.00; reactive control: BF01 = 1.93; Section S5). Thus, ACC sulcal morphology was neither associated with BI nor dependent on BI in its relationship to cognitive control in this sample.

## 4. Discussion

To our knowledge, this is the first study to investigate ACC sulcal patterns in individuals characterized for early BI. In this sample of adolescents, ACC sulcal asymmetry was associated with stronger proactive control (d’ context) on the AX-CPT, such that those with asymmetric patterns outperformed those with symmetric patterns. Given the sample size, these findings are preliminary. It is consistent with prior links between ACC sulcal morphology and cognitive control efficiency (Amiez et al., 2018; Cachia et al., 2014; Del Maschio et al., 2019; Huster et al., 2014; Tissier et al., 2018), and the direction converges with that work: asymmetric patterns were associated with lower Stroop interference in preschoolers (Cachia et al., 2014), and here with greater proactive control and less BX interference in adolescents. The present findings extend this work in two ways. First, previous studies relied on Stroop and Flanker paradigms, which index cognitive control efficiency but do not isolate control mode. The AX-CPT d’ context measure is designed specifically to capture proactive control, providing a more theoretically precise test. Second, ACC asymmetry was also associated with reduced BX interference, a marker of reactive control, extending the morphology-cognition link beyond proactive control to a distinct control mode within the same task.

After estimating both proactive (d′ context) and reactive control (BX interference) within the same model, ACC symmetry was associated with both measures, in opposite directions and to a similar degree (Supplementary Section S4). Two interpretations remain open. The pattern is consistent with ACC sulcal morphology relating to the balance between anticipatory and stimulus-driven control, as the dual mechanisms framework would suggest (Braver, 2012).

However, because d′ context and the BX probe interference effect are both derived, in part, from performance on BX trials and were strongly correlated in this sample, this pattern is equally consistent with morphology relating to a more general difference in control efficiency that is expressed on both measures. What the present data do indicate is that ACC sulcal morphology relates to both measures rather than to proactive control alone, and that the two associations are comparable in size.

ACC morphology was associated with cognitive control but not with BI, and Bayes factors moderately support no association rather than being underpowered (Section S5). The moderations provided weaker evidence, so it is inconclusive whether BI moderates the morphology-cognition association. Perhaps different aspects of ACC structure carry different kinds of information about development. Sylvester et al. (2016) found that cortical thickness continues to develop throughout the lifespan and may reflect the long-term effects of temperament and experience. They found that BI in early childhood predicted thinner dorsal ACC in adulthood. In contrast, ACC sulcal patterns measured in the present study are established prenatally and remain stable, so it cannot register later experience. It may therefore index early constraints on cognitive control efficiency that are orthogonal to temperament. BI may arise from partly distinct neurobiological pathways, such as amygdala reactivity and prefrontal-limbic connectivity, which may not be captured by ACC sulcal morphology. This interpretation is consistent with the findings from Cachia et al. (2014), who reported that ACC sulcal pattern predicted Stroop interference but not verbal working memory, suggesting some functional specificity in the cognitive processes associated with ACC folding. ACC sulcal morphology may be more closely linked to specific aspects of cognitive control rather than to broader cognitive or temperamental domains.

Exploratory analyses of whether 4-month infant reactivity moderated the association between ACC symmetry and cognitive control (Supplementary Section S3) did not yield a significant omnibus interaction for any control measure. Although the association between ACC symmetry and proactive control appeared strongest in the positive reactivity group, this within-group association did not survive correction for multiple comparisons and should therefore be interpreted with caution. Given the sample size, the study may have had limited power to detect such moderation effects, and whether early reactivity modulates the relationship between early neurodevelopmental constraints and later cognitive control remains an open question.

Because ACC sulcal patterns are established prenatally and remain stable across development (Cachia et al., 2021), the association between ACC asymmetry and cognitive control is consistent with the possibility that individual differences in proactive control are related to early-emerging brain organization rather than solely to later experience. This interpretation is necessarily tentative. The present design measured morphology and cognition at the same time and cannot therefore establish a causal developmental pathway; longitudinal work would be needed to test whether early sulcal morphology predicts the subsequent trajectory of cognitive control. Nevertheless, the present findings suggest that ACC sulcal morphology is associated with proactive control in adolescence, supporting further investigation as a potential early neurodevelopmental marker of cognitive control capacity.

There are several limitations to our study. First, our final analytic sample was modest (*N* = 62). Associations between brain structure and behavior are generally modest, and the present sample is small. In the ABCD study, MRI across modalities predicted approximately 20% of the variance in cognition (Pat et al., 2022), and statistically significant associations in small samples tend to overstate the true effect. The associations reported here should thus be treated as preliminary and in need of replication in larger samples. This study’s modest sample size also may limit generalizability and likely reduced statistical power to detect interaction effects in our exploratory moderation analyses. Second, proactive and reactive control were operationalized using a single task, the AX-CPT. In this task, d’ context and BX interference both rely in part on performance on BX trials, which may contribute to their strong negative association and explain why it is difficult to estimate how independent they are from each other based on behavior.

Confirming these associations with other paradigms that distinguish between the two control modes, such as cued or proportion-congruency variants of the Stroop and Flanker tasks, would strengthen confidence that the observed associations reflect differences in control mode rather than task-specific demands. Third, the current study examined cognitive control at only a single time point, precluding examination of how ACC morphology relates to developmental trajectories of cognitive control. Finally, our small temperament subgroups and our sample being predominantly White further limited external validity in this study.

Future studies with larger samples will be needed to adequately power tests of whether early temperament moderates the association between ACC sulcal morphology and cognitive control. Functional MRI could also help determine whether ACC morphology is associated with differential ACC recruitment during proactive and reactive control on the AX-CPT. Longitudinal designs could track developmental trajectories of proactive and reactive control as a function of ACC morphology, while longitudinal mediation analysis could test whether ACC morphology is indirectly associated with later anxiety risk through individual differences in cognitive control.

In this sample of adolescents, ACC sulcal asymmetry was associated with more proactive and less reactive cognitive control, with associations that did not differ across the two control modes. These preliminary findings extend the literature on ACC morphology and cognitive control by distinguishing between proactive and reactive modes of control and by examining these associations in a sample characterized for early temperament. The findings that ACC morphology was associated with cognitive control but not with BI further suggest it may index early neurodevelopmental constraints on cognitive control rather than on temperament per se. Replication in larger samples and in longitudinal designs will be needed to establish whether early neurodevelopmental variation contributes to BI-related mental health trajectories.

