## Supplemental Materials for "ACC Sulcal Morphology as a Neurodevelopmental Marker of Proactive and Reactive Cognitive Control among Adolescents"

This research was supported by the National Institute of Mental Health, Grant/Award Number: K23MH130751 (to E. A. V.), U01MH093349 (to N. A. F.), and Intramural Research Project ZIAMH002782 (to D. S. P.). Hence, this research was supported [in part] by the Intramural Research Program of the National Institutes of Health (NIH). The contributions of the NIH author(s) are considered Works of the United States Government. The findings and conclusions presented in this paper are those of the author(s) and do not necessarily reflect the views of the NIH or the U.S. Department of Health and Human Services.

### Contents

Supplementary Section S1. AX-CPT Task Parameters and Trial-Level Exclusions

Supplementary Section S2. Left x Right Architecture Analyses

Supplementary Section S3. Infant Reactivity Moderation

Supplementary Section S4. Joint Proactive-Reactive Model

### Supplement

#### Supplementary Section S1. AX-CPT Task Parameters and Trial-Level Exclusions

Proactive and reactive control were assessed at the 15-year assessment using the AX-Continuous Performance Test (AX-CPT; Cohen et al., 1999), in which a cue is followed by an "X" probe. The task had 316 trials across 4 blocks, with trial proportions of 55% AX, 15% AY, 15% BX, and 15% BY, across 319 total trials across 4 blocks. The cue-probe delay was randomized between 1,400 and 1,600 ms. Cue letters were presented in cyan and probe letters in white on a black background, in 60-point Courier New font. Stimuli were presented using E-Prime 2.0.

Trials with reaction times more than 3 standard deviations above or below each participant's mean were excluded (less than 3% of trials). Participants were required to achieve at least 60% accuracy on BY trials for inclusion in analyses. Of the 66 participants with valid ACC sulcal classification, 3 did not complete the 15-year AX-CPT and 1 completed fewer than 1 task block (fewer than 80 of the 319 trials, under 25% of the task). These participants were excluded from cognitive control analyses, yielding a final analytic sample of 62.

#### Supplementary Section S2. Left x Right Architecture Analyses

In the main analyses, ACC morphology was operationalized as sulcal symmetry. As a robustness check, we repeated the behavioral inhibition analysis operationalizing morphology as the interaction between left- and right-hemisphere sulcal architecture (single vs. double parallel type), reflecting the original analysis approach. Neither left-hemisphere architecture ($F(1, 55) = 0.02$, $p = .893$), right hemisphere architecture ($F(1, 55) = 0.99$, $p = .325$), nor their interaction ($F(1, 55) = 0.15$, $p = .704$) predicted BI ($N = 62$), consistent with the symmetry-based analysis reported in the main text. ACC morphology was not associated with BI under either operationalization.

#### Supplementary Section S3. Infant Reactivity Moderation

To examine whether 4-month infant reactivity groups moderated the association between ACC sulcal symmetry and cognitive control, we added an ACC asymmetry by reactivity group interaction to each model. For proactive control, (d' context), the omnibus interaction was not significant ($F(2, 50) = 1.18$, $p = .315$), nor was the main effect of the reactivity group ($F(2, 50) = 0.16$, $p = .849$. Planned within-group comparisons using estimated marginal means (FDR-corrected across groups) indicated that the symmetry effect on d' context was largest in the positive reactivity group (symmetric $M = 2.80$, $SE = 0.23$ vs. asymmetric $M = 3.65$, $SE = 0.28$; $b = 0.85$, $SE = 0.35$, $t(50) = 2.44$, $p$FDR = .054), at a trend level, with no effect in the negative ($p$FDR = .239) or control ($p$FDR = .873) groups.

The interaction was also non-significant for both reactive control measures (BX interference error rate: $F(2, 50) = 0.26$, $p = .774$; reaction time: $F(2, 50) = 1.31$, $p = .278$). Within-group comparisons yielded no effects surviving FDR correction for either measure (error rate: control *p*FDR = .445, negative *p*FDR = .068, positive *p*FDR = .157; reaction time: control *p*FDR = .804, negative *p*FDR = .055, positive *p*FDR = .557). Given the non-significant omnibus interactions, these within-group patterns are exploratory and should be interpreted with caution.

**Supplement Section S4. Joint proactive-reactive model**

To compare the magnitude of the ACC symmetry effect across the two control modes, d' context and the BX probe interference error rate were regressed simultaneously on ACC symmetry and covariates (gender, age at scan, behavioral inhibition, race/ethnicity, and 4-month reactivity group) in a single structural equation model (SEM) (lavaan package) estimated with maximum likelihood and full-information maximum likelihood (fiml) for missing data. The residual covariance between the two outcomes was freely estimated, so that each symmetry effect is interpretable net of the what the two variables share. Both outcomes were standardized and retained their original scoring, in which higher d' context indication more proactive control and higher BX interference means more reactive control use. Effects of equal size on the two outcomes therefore appear as coefficient of equal magnitude and opposite sign, and this was tested as the sum of the two coefficients.

Symmetric patterns were associated with both lower proactive control (*d'* context; $b$ = -0.61, $p$ = .018) and greater BX interference ($b$ = +0.75, $p$ = .001). The sum of the two coefficients did not differ from zero (0.14, $z$ = 0.57, $p$ = .569), indicating that the two effects did not differ in magnitude. D' context and BX interference were negatively associated after accounting for all predictors (standardized residual covariance = 0.61, $p$ <.001). With the reaction-time reactive measure, the pattern was in the same direction but weaker (symmetric $b$ = +0.52, $p$ = .067; residual covariance = −0.11, $p$ = .388).

The joint model was saturated ($df$ = 0); therefore, the global fit indices were not informative and are not reported. Given the small sample size, these estimates are preliminary.

**Supplementary Section S5. Bayes Factor for the Behavioral Inhibition Analyses**

Conventional null hypothesis testing cannot distinguish evidence for the absence of an association from a statistically underpowered dataset. To address this, Bayes factors were computed for the behavioral inhibition analyses using the BayesFactor package (Morey & Rouder, 2024) in R. for each test, a model including the effect of BI was compared with an identical model omitting the variable of interest with the same covariates (gender, race/ethnicity and age at scan) for each model. Default prior scales included in the package were left unchanged (Cauchy prior on standardized effect sizes, r = 0.5 for categorial predictors and r = 0.354 for continuous covariates) and analyses were restricted to complete cases.

Bayes factors are reported as $BF_{01}$, evidence for the null hypothesis (no association with BI) relative to the alternate hypothesis, such that values above 1 favor no association. Following conventional benchmarks, values between 1 and 3 are considered anecdotal, values greater than 3, below 10 is moderate and values above 10 provide strong evidence (Lee & Wagenmakers, 2013).

For the association between ACC sulcal symmetry and BI, the model without symmetry provided moderate evidence for no association between the two ($BF_{01}$ = 3.79, $N$ = 62). For the symmetry-by-BI interactions, Bayes factors were close to 1 for both proactive control ($BF_{01}$ = 2.00, $N$ = 58) and reactive control ($BF_{01}$ = 1.93, $N$ = 58), which is considered inconclusive and may mean that the data was underpowered rather than supporting that there is no moderation. These moderation results should therefore be interpreted carefully as they are not evidence against an interaction.
